# Neural mechanisms for reward-modulated vector learning and navigation: from social insects to embodied agents

**DOI:** 10.1101/045559

**Authors:** Dennis Goldschmidt, Poramate Manoonpong, Sakyasingha Dasgupta

## Abstract

Despite their small size, insect brains are able to produce robust and efficient navigation in complex environments. Specifically in social insects, such as ants and bees, these navigational capabilities are guided by orientation directing vectors generated by a process called path integration. During this process, they integrate compass and odometric cues to estimate their current location as a vector, called home vector for guiding them back home on a straight path. They further acquire and retrieve path integration-based vector memories anchored globally to the nest or visual landmarks. Although existing computational models reproduced similar behaviors, they largely neglected evidence for possible neural substrates underlying the generated behavior. Therefore, we present here a model of neural mechanisms in a modular closed-loop control - enabling vector navigation in embodied agents. The model consists of a path integration mechanism, reward-modulated global and local vector learning, random search, and action selection. The path integration mechanism integrates compass and odometric cues to compute a vectorial representation of the agent’s current location as neural activity patterns in circular arrays. A reward-modulated learning rule enables the acquisition of vector memories by associating the local food reward with the path integration state. A motor output is computed based on the combination of vector memories and random exploration. In simulation, we show that the neural mechanisms enable robust homing and localization, even in the presence of external sensory noise. The proposed learning rules lead to goal-directed navigation and route formation performed under realistic conditions. This provides an explanation for, how view-based navigational strategies are guided by path integration. Consequently, we provide a novel approach for vector learning and navigation in a simulated embodied agent linking behavioral observations to their possible underlying neural substrates.

**Author Summary:** Desert ants survive under harsh conditions by foraging for food in temperatures over 60° C. In this extreme environment, they cannot, like other ants, use pheromones to track their long-distance journeys back to their nests. Instead they apply a computation called path integration, which involves integrating skylight compass and odometric stimuli to estimate its current position. Path integration is not only used to return safely to their nests, but also helps in learning so-called vector memories. Such memories are sufficient to produce goal-directed and landmark-guided navigation in social insects. How can small insect brains generate such complex behaviors? Computational models are often useful for studying behavior and their underlying control mechanisms. Here we present a novel computational framework for the acquisition and expression of vector memories based on path integration. It consists of multiple neural networks and a reward-based learning rule, where vectors are represented by the activity patterns of circular arrays. Our model not only reproduces goal-directed navigation and route formation in a simulated agent, but also offers predictions about neural implementations. Taken together, we believe that it demonstrates the first complete model of vector-guided navigation linking observed behaviors of navigating social insects to their possible underlying neural mechanisms.

## Introduction

Social insects, including ants and bees, have evolved remarkable behavioral capabilities for navigating in complex dynamic environments, which enable them to survive by finding vital locations (e.g., food sources). For example, desert ants are able to forage and find small, sparsely distributed food items in a featureless environment, and form stereotyped and efficient routes between their nest and reliable food sources [1–4]. These navigational behaviors not only rely on sensory information, mainly from visual cues, but also on internal memories acquired through learning mechanisms [5]. Such learned memories have shown to be based on orientation directing vectors, which are generated by a process called path integration (PI) [6].

### Vector navigation in social insects

In PI, animals integrate angular and linear ego-motion cues over time to produce an estimate of their current location with respect to their starting point. This vector representation is called home vector (HV) and is used by social insects to return back to the home on a straight path. Many animals have been shown to apply PI, including vertebrate [7] and invertebrate species [8]. While PI has mainly been observed in homing behavior, it can also serve as a scaffold for spatial learning of food sources [9]. This learning ability is apparent in the waggle dance of honey bees [10, 11], in which the distance and direction to a goal are encoded by the duration and direction of the dance, respectively. The vector representation of food sources is called global vector (GV), because it connects the food location to the nest which is the global origin of PI. After returning from a successful foraging run, insects re-apply this vector information in subsequent foraging runs [12, 13].

Although PI plays a key role in navigating through environments where visual cues, such as landmarks, are abundant, it also influences navigational behaviors in cluttered environments [14]. If an ant follows a learned GV repeatedly, it learns the heading directions at local landmarks along the path [15]. These heading directions are view-based from the visual panorama surrounding the ant [16, 17], and vector-based with additional information about the path segment length [15, 18]. The latter vector memories are also termed local vectors (LV), because their origin is linked to local landmarks instead of the nest [19]. Besides spatial learning of locations and routes, searching patterns of desert ants have also shown to be influenced by PI [20, 21]. In addition, it has been observed that after directionally biased outbound walks, desert ants return back to the nest with systematic errors, in which the animal misestimates the home position by a short distance right in front of the actual nest [22]. These errors have been observed in several species in vertebrates and invertebrates [7].

### Neural substrates of social insect navigation

Neural substrates of social insect navigation have yet to be completely identified, but previous findings of neural representations of compass cues and visual sceneries may provide essential information about how PI and vector learning is achieved in neural systems [23–26]. In particular, neurons in the central complex, a protocerebral neuropil in the insect brain, have shown to be involved in visually guided navigation. The main sensory cue for PI in social insects is derived from the linear polarization of scattered sunlight [27–29]. Specialized photoreceptors in the outer dorsal part of the insect eye detect certain orientations of linear polarization, which depend on the azimuthal position of the sun. A distinct neural pathway processes polarization-derived signals leading to neurons in the central complex, which encode azimuthal directions of the sun [30]. In a recent study, Seelig and Jayaraman [23] placed a fruit fly tethered on a track ball setup in a virtual environment and measured the activity of neurons in the central complex. They demonstrated that certain neurons in the ellipsoid body, which is a toroidal subset in the central complex, encode for the animal’s body orientation based on visual landmarks and angular self-motion. When both visual and self-motion cues are absent, this representation is maintained through persistent activity, which is a potential neural substrate for short-term memory in insects [31]. A similar neural code of orientations has been found in the rat limbic system [32]. These so-called head direction (HD) cells are derived from motor and vestibular sensory information by integrating head movements through space. Thus, neural substrates of allothetic compass cues have been found in both invertebrate and vertebrate species. These cues provide input signals for a potential PI mechanism based on the accumulation of azimuthal directions of the moving animal as previously proposed by Kubie and Fenton [33].

### Computational models of vector-guided navigation

Because spatial navigation is a central task of biological as well as artificial agents, many studies have focused on computational modeling of such behavioral capabilities (see [34] for review). Computational modeling has been invaluably successful in exploring the link between neural structures and their behavioral function. It allows for hypotheses about the underlying mechanisms to be defined precisely and their generated behavior can be examined and validated qualitatively and quantitatively with respect to experimental data.

Most models of PI favor a particular coordinate system (Cartesian or polar) and reference frame (geo-or egocentric) to perform PI based on theoretical and biological arguments [35]. While some models [22, 36] include behavioral data from navigating animals in order to argue for their proposed PI method, others [37–39] have applied neural network models to investigate possible memory mechanisms for PI. Despite the wide variety of models, none of these models have been implemented on realistic embodied artificial agents in order to provide results of their model in the ecological context of animals. Furthermore, possible links between PI and navigational capabilities, including spatial learning and memory, have largely been ignored. Kubie and Fenton [33] proposed a PI model based on the summation of path segments with HD accumulator cells, which are individually tuned to different HDs and hypothesized to encode how far the animal travelled in this direction. These summated path vectors are then stored in a fixed memory structure called shortcut matrix, which is used for navigating towards goals. Although this model is based on HD cells and therefore presented as for mammalian navigation, recent findings in Drosophila [23] demonstrate that similar HD accumulator cells can also be hypothesized for insect navigation. Similar HD accumulator models have been applied for chemo-visual robotic navigation [40] and PI-based homing behavior [39]. Cruse and Wehner [41] presented a decentralized memory model of insect vector navigation to demonstrate that the observed navigational capabilities do not require a map-like memory representation. Their model is a cybernetical network structure, which mainly consists of a PI system, multiple memory banks and internal motivational states that control the steering angle of a simulated point agent. The PI system provides the position of the agent given by euclidean coordinates, which are stored as discrete vector memories when the agent finds a food location. To our knowledge, this model is the first and only modeling approach which accounts for behavioral aspects of insect vector navigation. However, although they introduce a learning rule for so-called quality values of stored vectors in a more recent version of the model [42], their model does not account for how the navigation vectors are represented and learned in a neural implementation.

### Our approach

Based on these findings, in this paper, we present a novel model framework for PI and adaptive vector navigation as observed in social insects. The framework is applied as closed-loop control to an embodied agent and consists of five functional subparts: 1) a neural PI mechanism, reward-modulated learning rules for 2) GV and 3) LV memories, 4) random search, and 5) an adaptive action selection mechanism. Using this framework on a simulated embodied agent, we demonstrate that we can not only reproduce qualitatively similar behavior as observed in social insects, but further predict experimental data from behavioral and neurobiological studies.

Based on population encoding of heading directions in circular arrays, we apply PI by accumulating speed-modulated HD signals through a self-recurrent loop. The final HV representation is computed by local excitation-lateral inhibition connections, which projects accumulated heading directions onto the array of output neurons. The activity of these neurons encodes the vector angle as the position of maximum firing in the array, and the vector length as the amplitude of the maximum firing rate in the array.

The self-localization ability of PI allows social insects to learn places and supports the formation of visually guided routes. We apply a reward-modulated associative learning rule [43–45] to learn vector representations based on PI. Two types of vector representations are learned in two seperate neural circuits, the so-called GV and LV. A GV connects the nest to a rewarding food location. GVs are learned by the three-factored association of the PI state, a context-dependent state, and the respective reward received at the food location. This association induces weight changes in plastic synapses connecting the context-dependent unit to the so-called GV array, which represents the GV. The context-dependent unit activates the GV representation in the array, and therefore represents a motivational state for goal-directed foraging. Using the GV learning rule, the agent is able to learn rewarding locations and demonstrate goal-directed navigation. Because of the substractive weighting of GV and HV in the action selection mechanism, it can compensate for unexpected detours from the original trajectory, such as obstacles.

LVs are vectors, which originate from prominent locations, such as landmarks or panoramic views [19]. In our model, LVs are learned in a similar form as the GVs, but here by the association of a reference PI state, a time-decaying landmark detection state (context), and the reward. This changes synaptic weights in the so-called LV array. The inputs to this array are several units which encode for the time-decaying landmark detection as an eligibility trace. Only one of these units is active at each time, which is the most recently detected landmark. We show that the LV learning rule leads to learning LV in a recursive way from the reward location to the nest. To ensure that the agent learns LVs from one landmark to another, we introduce a value system which encodes how successful a landmark is in leading to the goal. These so-called LV values increase by correlations of the eligibility traces with the reward and act as additional reward signals in the learning rules.

Taken together, our model is a novel framework for generating and examining social insect navigation based on PI and vector representations. It is based on neural mechanisms, which are related to neurobiological findings in the insect central complex. Therefore, we provide a computational approach for linking behavioral observations to their possible underlying neural substrates. In the next section, we will describe the proposed model for reward-modulated vector learning and navigation. The results section will provide detailed descriptions of our experimental setups and simulation results. Finally, conclusions and implications of our model with respect to behavioral and neurobiological studies are discussed in the last section.

## Model

In this paper we propose an insect-inspired model of vector-guided navigation in embodied agents using modular closed-loop control. The model (see Fig. 1A) consists of five parts: 1) a neural PI mechanism, plastic neural circuits for reward-based learning of 2) GV and 3) LV memories, 4) random search, and 5) action selection. The neural mechanisms in our model receive multimodal sensory inputs from exteroceptive and proprioceptive sensors to produce a directional signal based on a vector. This vector is represented by the activity of circular arrays, where the position of the maximum indicates its direction and the amplitude at this position indicates its length.

**Figure 1:**
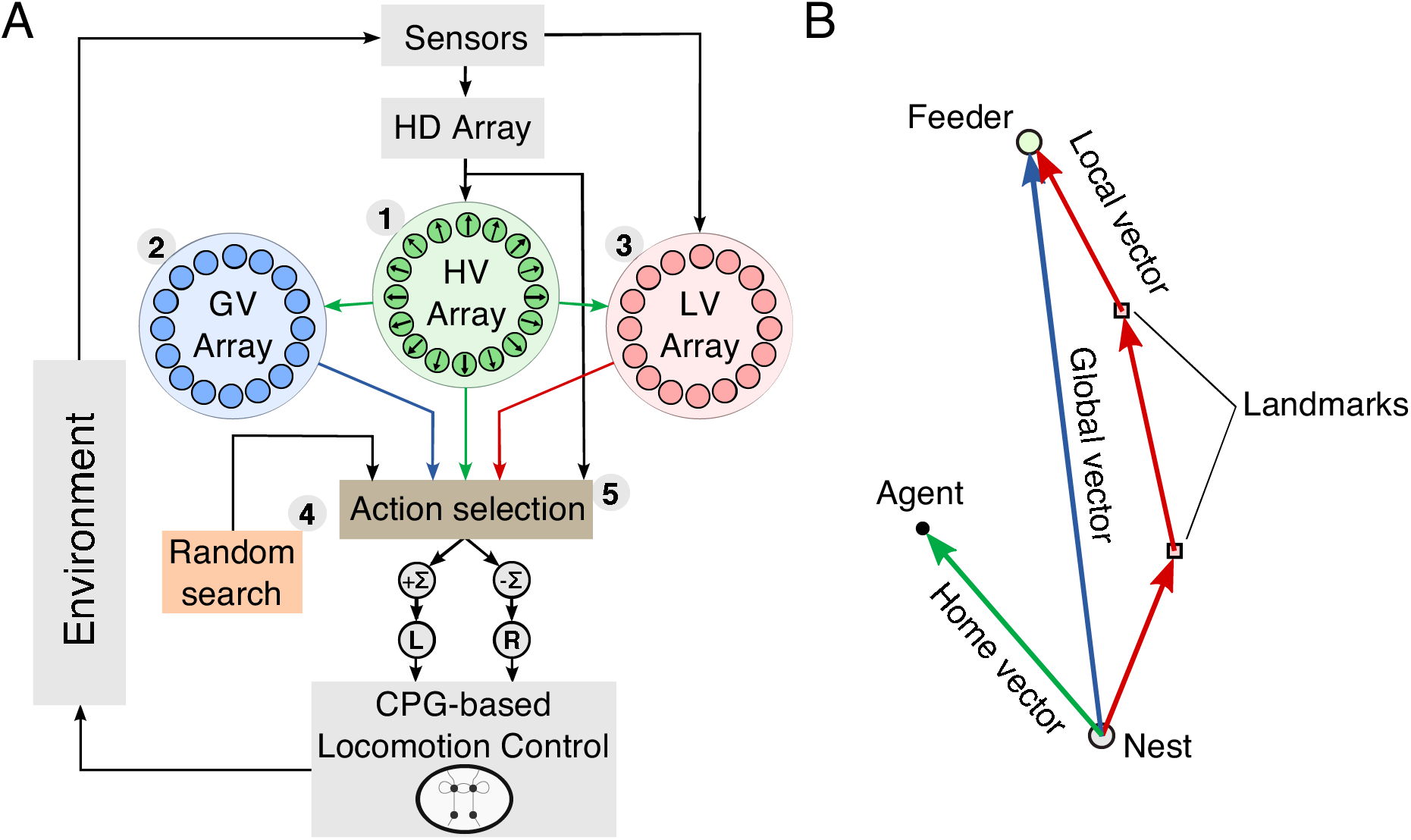
Schematic diagram of the modular closed-loop control for vector navigation. A) The model consists of a neural path integration (PI) mechanism (1), reward-modulated global (2) and local vector (3) learning, random search (4), and an adaptive action selection mechanism (5). Vector information for guiding navigation is computed and represented in the activity of circular arrays. The home vector (HV) array is the output of the PI mechanism and is applied for homing behavior and as a scaffold for global (GV) and local vector (LV) learning. These three vector representations and random search are integrated through an adaptive action selection mechanism, which produces the steering command to the CPG-based locomotion control. B) Spatial representation of the different vectors used for navigation. The HV is computed by PI and gives an estimate for the current location of the agent. In general, GVs connect the nest to a rewarding location. LVs are anchored in rewarding locations other than the nest, such as prominent landmarks.

The PI mechanism integrates angular and linear motion cues to generate the HV, which is an estimate connecting the nest with the current location of the agent. Therefore, social insects mainly use it to return back to the nest on a straight path [6]. In addition, this vector has been shown to be used as a scaffold for spatial learning of rewarding goals [46]. Here we apply two types of learned vectors to guide navigation of the agent. A GV is learned by association of the path integration state with the received reward, and therefore links the nest to a rewarding location. LVs are defined by originating from stationary landmarks to rewarding locations (see Fig. 1B). Details about the applied learning rules will be described below. The three vector representations (HV, GV, LV) and random search are integrated through an adaptive action selection mechanism allowing exploration as well as the exploitation and competition of learned vectors. We evaluate our model using a two-dimensional point agent as well as a hexapod walking robot (see S1 Text for details).

### Path integration (PI) mechanism for home vector (HV) representation

The PI mechanism (Fig. 2) is a multilayered neural network consisting of circular arrays, where the final layer’s activity pattern represents the HV. Neural activities of the circular arrays represent population-coded compass information and rate-coded linear displacements. Incoming signals are sustained through leaky neural integrator circuits, and they compute the HV by local excitatory-lateral inhibitory interactions.

**Figure 2:**
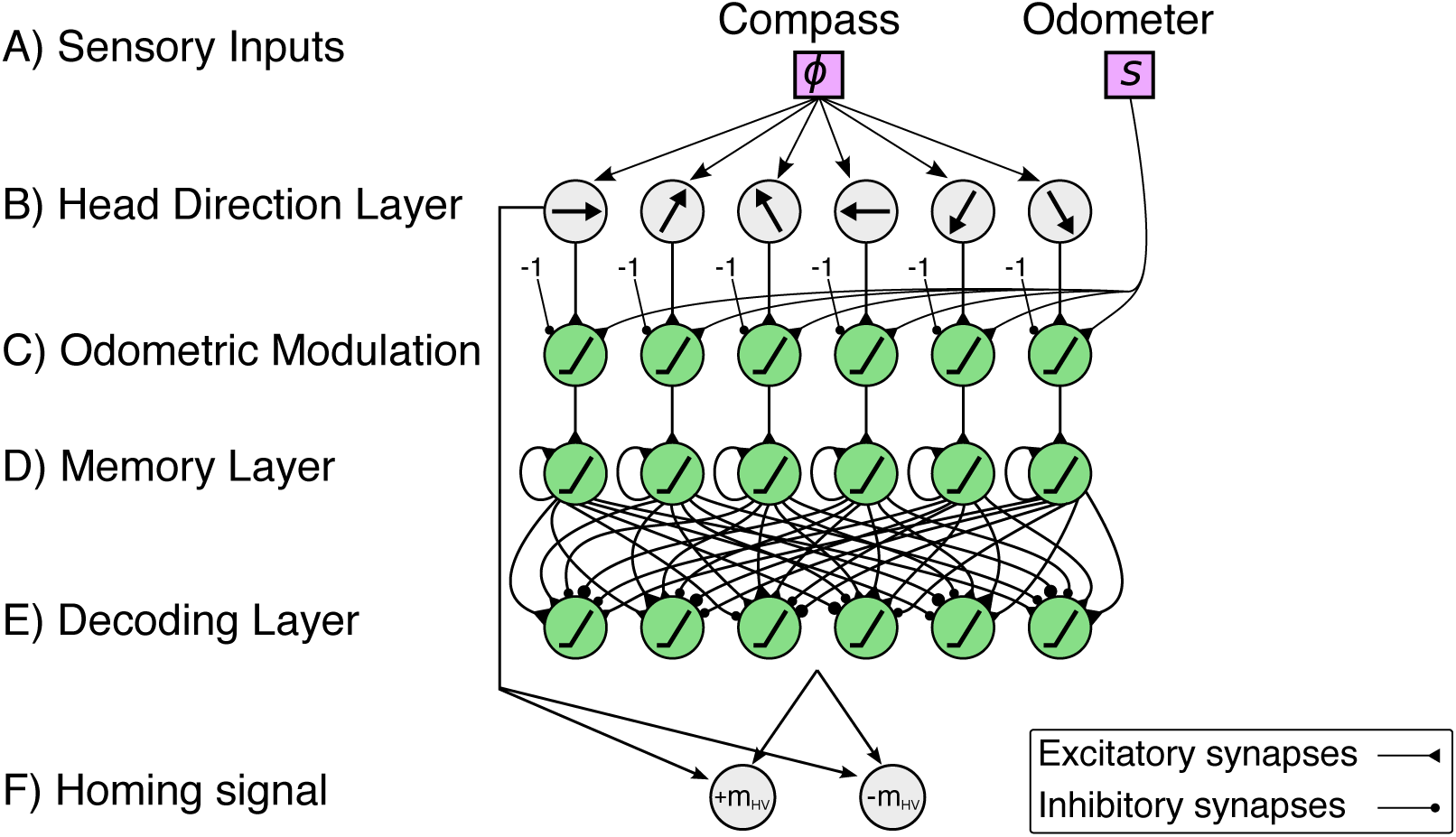
Multilayered neural network of the proposed path integration (PI) mechanism. A) Sensory inputs from a compass sensor (*ϕ*) and odometer (*s*) are provided to the mechanism. B) Neurons in the head direction (HD) layer encodes the sensory input from a compass sensor using a cosine response function. Each neuron encodes a particular preferred direction enclosing the full range of 2*π*. Note that the figure depicts only six neurons for simplicity. C) An odometric sensory signal (i.e., walking speed) is used to modulate the HD signals. D) The memory layer accumulates the signals by self-recurrent connections. E) Cosine weight kernels decode the accumulated directions to compute the output activity representing the home vector (HV). F) The difference between the HV angle and current heading angle is used to compute the homing signal.

#### A) Sensory inputs

The PI mechanism receives angular and linear motion cues as sensory inputs. Like in social insects, angular cues are derived from allothetic compass cues. We employ a compass sensor which measures the angle *ϕ* of the agent’s orientation. In insects, this information is derived from the combination of sun‐ and skylight compass information [6]. Odometry measures the walking speed *s* of the agent. For the embodied agent employed here (i.e., a hexapod robot), the walking speed is computed by accumulating steps and averaging over a certain time window. These step counting signals are derived from the motor signals. The input signals have value ranges of

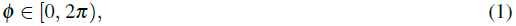

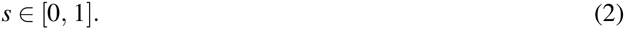

#### B) Head direction layer

The first layer of the neural network is composed of HD cells with activation functions

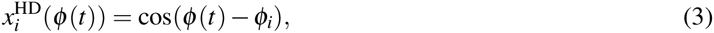

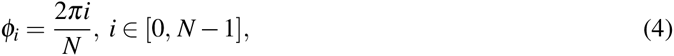

where the compass signal *ϕ*(*t*) is encoded by a cosine response function with *N* preferred directions *ϕ*_*i*_, ϵ [0,2*π*). The resolution is determined by 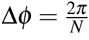 and the coarse encoding of variables, here angles, by cosine responses allows for high accuracy and optimized information transfer [47]. Coarse coding has been shown to be present in different sensory processing in the insect brain, including olfactory [48] and visual processing [49].

#### C) Odometric modulation of head direction signals

The second layer acts as a gating mechanism (G), which modulates the neural activity using the odometry signal *s* (ϵ [0, 1]). Therefore, it encodes in its activit,the travelled distances of the agent. The gating layer units decrease the HD activities by a constant bias of 1, so that the maximum activity is equal to zero. A positive speed increases the signal linearly. The gating activity is defined as follows:

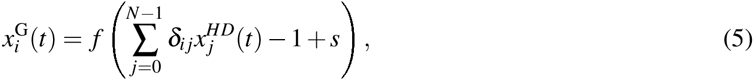

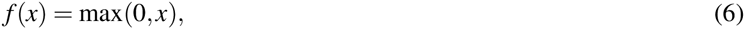

where *δ*_*ij*_ is the Kronecker delta, i.e., first layer neurons *j* and second layer neurons *i* are connected one-to-one. Examples of different speed-modulated gating activities are shown in Fig. 3.

Figure 3: Example of speed modulation using a gating mechanism with 36 neurons (second layer, see Fig. 2C). The agent’s heading direction is *ϕ* = 135° and the speed s is set to (a) 0.25, (b) 0.5, and (c) 1.0.

#### D) Memory layer

The third layer is the so-called memory layer (M), where the speed-modulated HD activations are temporally accumulated through self-excitatory connections:

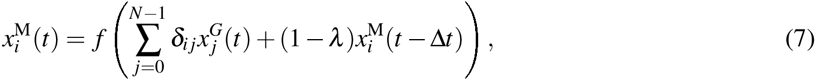

where *λ* is a positive constant defined as the integrator leaking rate, which indicates the loss of information over time.

#### E) Decoding layer

The final and fourth layer decodes the activations from the memory layer to produce a vector representation, i.e., the HV, which serves as the output of the mechanism referred to as PI state:

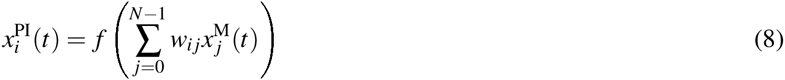

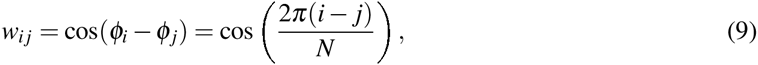

where *w*_*ij*_ is a cosine kernel, which decomposes the projections of memory layer actitivities of the *j*th neuron to the *i*th neuron’s preferred orientation. The resulting HV is encoded by the average position of maximum firing in the array (angle *θ*_*HV*_) and the sum of all firing rates of the array (length *l*_*HV*_). We calculate the position of maximum firing using the population vector average given by

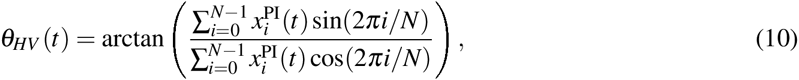

where the denominator is the *x* coordinate of the population vector average, and the numerator is the *y* coordinate. See Fig. 4 for example output activities of the decoding layer neurons.

**Figure 4:**
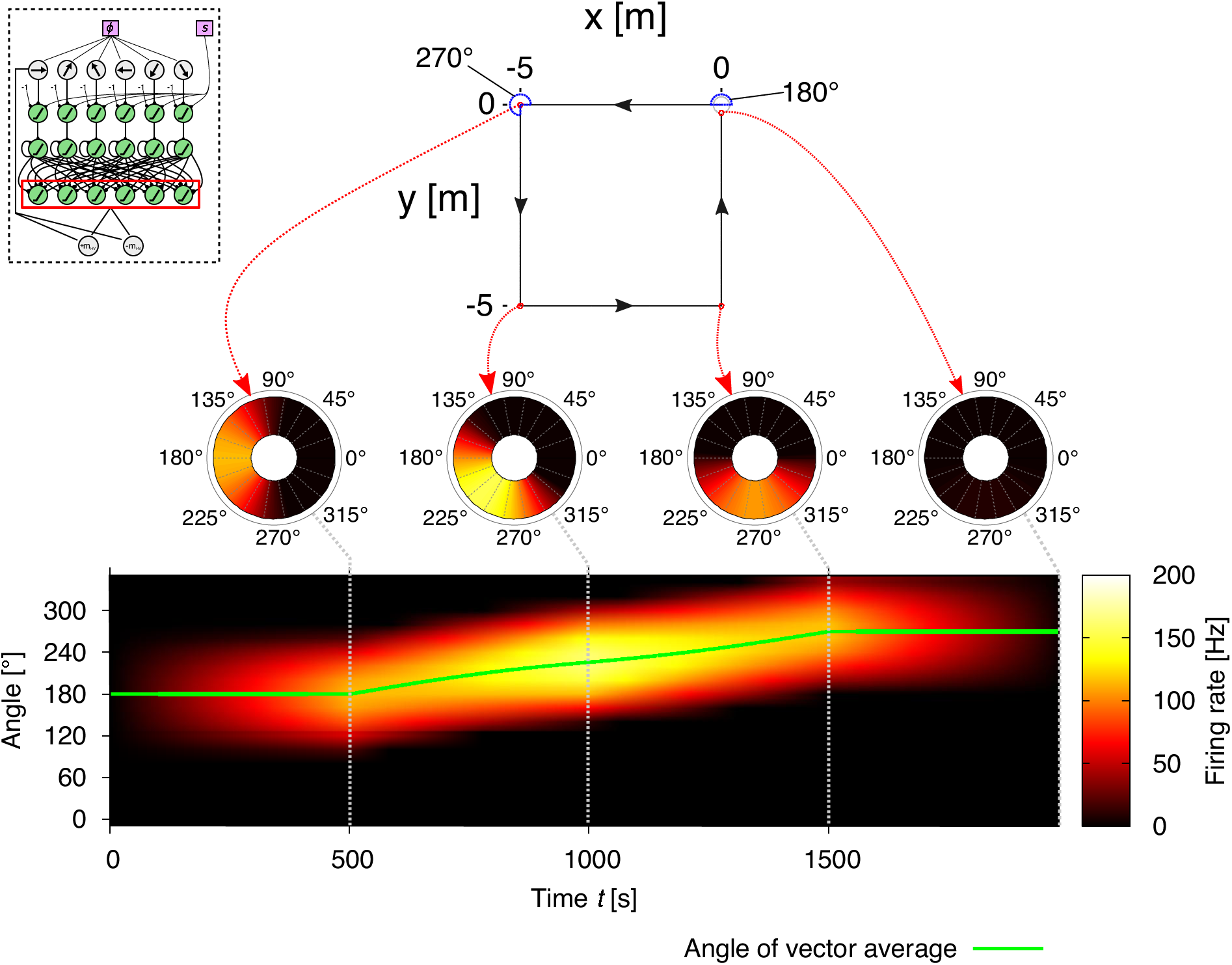
Example of vector representations based on the neural activities of the decoding layer (seeFig. 2E) in the path integration (PI) mechanism for a square trajectory. The agent runs for 5 m in one of the four directions (180°, 270°, 0°, 90°), thus finally returning to the starting point of its journey. The coarse encoding of heading orientations lead to a correct decoding of memory layer activities. Thus, the output activities of the PI mechanism represent the home vector (HV), where the position of the maximum firing rate is the angle and the amplitude of the maximum firing rate is the length of the vector. Note that, as the agent returns to the home position, the output activities are suppressed to zero resulting from the elimination of opposite directions.

#### F) Homing signal

To apply the HV for homing behavior, i.e. returning home on a straight path, the vector is inverted by a 180^°^ rotation. The difference between the heading direction *ϕ* and the inverted HV direction *θ*_HV_ – *π* is used for steering the agent towards home. The agent applies homing by sine error compensation, which defines the motor command

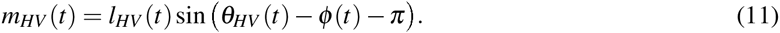

This leads to right (*m*_*HV*_ < 0) and left turns (*m*_*HV*_ > 0) for negative and positive differences, respectively, and thereby decreasing the net error at each step. The underlying dynamical behavior of this sine error compensation is defined by a stable and an unstable fixed point (see S2 Text). This leads to dense searching behavior around a desired position, where the error changes rapidly [35].

### A reward-modulated learning rule for acquiring and retrieving vector memories

We propose a heterosynaptic, reward-modulated learning rule [43–45] with a canonical form to learn GVs and LVs based on four factors (see Fig. 5): a context-dependent state, an input-dependent PI state, a modulatory reward signal, and the vector array state.

**Figure 5:**
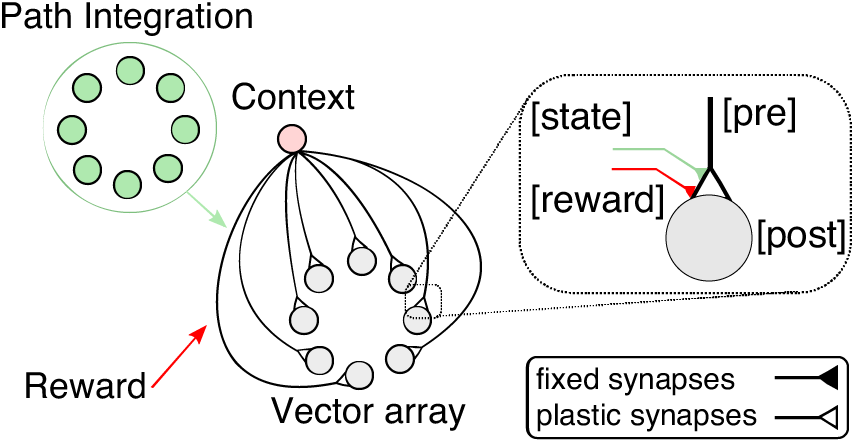
Canonical vector learning rule involves associations of path integration (PI) states with contextdependent and reward signals. Both, global (GV) and local vector (LV) memories are acquired and expressed by this learning circuit. For GVs, we associate the home vector (HV) array activities with the food reward given an active foraging state (outward journey). LVs are learned through association of a reference vector with the local reward (food or rewarding landmark) given a detected landmark. For details, see text below.

As in the PI mechanism, GVs and LVs are computed and represented in circular arrays. The context-dependent state activates the vector representation, and thus retrieves the vector memory. The association between the PI-based state and the reward signal modulates the plastic synapses connecting the context unit (presynaptic) with the vector array units (postsynaptic).

### Global vector (GV) learning

We propose a reward-modulated associative learning rule for spatial learning of GVs. A GV is learned through the association of the PI state, which represents the conditioned stimulus, and of the reward signal, which represents the unconditioned stimulus. The associated information is used by the agent on future foraging trips to steer towards the rewarding location. The received reward is an internally generated signal based on food reward due to visiting the feeder.

For GVs, the context-dependent unit (see Fig. 5) is a unit that represents the agent’s foraging state, i.e., inward or outward. Here we apply a simple binary unit given by

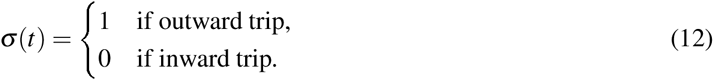

The context-dependent unit projects plastic synapses onto a circular array that represents the GV. The GV array has the same number of neurons, thus the same preferred orientations as the PI array. In this way, each neuron *i* ϵ [0,*N* – 1] has a preferred orientation of 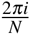. The activity 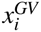 of the GV array is given by

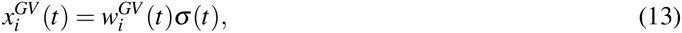

where 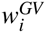 are the weights of the plastic synapses. For these synapses, we apply a reward-modulated associative learning rule given by

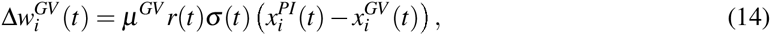

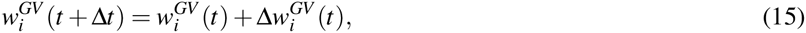

where *μ*^*GV*^ is the learning rate, and 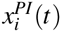 is the PI activity in the direction 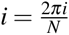.The food reward *r*(*t*) at the feeder is given by

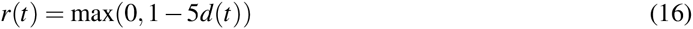

where *d*(*t*) is the agent’s distance to the feeder. Due to the delta rule-like term 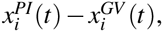, the weights 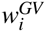 approach same values as the activities of the PI state at the rewarding location. Thus, the weights represent the static GV to the rewarding location (feeder). After returning back home, the agent applies the angle *θ*_*GV*_ of the GV to navigate towards the feeder using error compensation. The motor signal of the GV

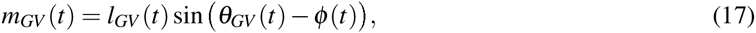

is applied together with the homing signal *m*_*HV*_ and random search *m*_*ε*_, where *l*_*GV*_ is the length of the GV. Random search is a noise signal drawn from a Gaussian distribution *N* (mean, S.D.):

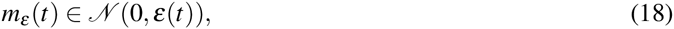

with an adaptive exploration rate ε(*t*) given by

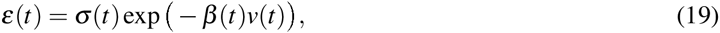

where *v*(*t*) is an estimate for the average food reward received over time and *β*(*t*) is the inverse temperature parameter. We define *v* by the recursive formula

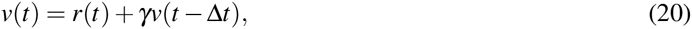

where *v*(*t*) is a lowpass filtered signal of the received food reward *r*(*t*) with discount factor *γ* = 0.995. Convergence of goal-directed behavior is achieved for *ε* below a critical value, which depends on the choice of *β*. We assume that *ε* and *v* are based on a probability distribution with fixed mean. We derive a gradient rule, which leads to minimization of the Kullback-Leibler divergence between the distribution of *ε*(*v*) and an optimal exponential distribution (see S3 Text for a derivation). The learning rule is given by

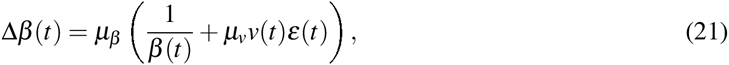

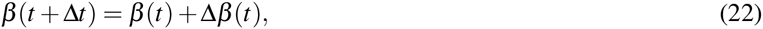

where *μ*_β_ = 10^‒6^ is a global learning rate, *μ*_*v*_ = 10^2^ is a reward-based learning rate. The adaptation of beta is characterized by small changes scaling with the square root of time, while the term containing *v*(*t*) allows for exploitation of explored food rewards to further decrease *ε* through *β*. In ecological terms, such exploitation of sparse distributed resources is crucial for the survival of an individual as well as the whole colony [20, 50, 51].

The final motor command ∑ for goal-directed navigation is given by the linear combination

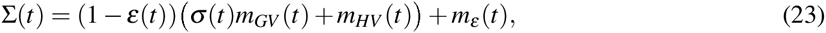

where outward trips are controlled by the balance of random walk and global-vector guided navigation depending on the exploration rate *ε*, while inward trips are controlled solely by the homing signal *m*_*HV*_. The combination of the two sinusoidals is equivalent to a phase vector (phasor) addition resulting in a phasor, which connects the current position of the agent with the learned feeder location (see S2 Text for a derivation).

### Local vector (LV) learning

PI is a computation, in which errors are accumulated continuously with each update step [35, 52]. The effect of noise, which is abundant in real-life systems, therefore also influences the learned representations of the GV. It has been shown that the PI error scales with the square root of distance to the nest. The same relation holds for GV representations, which therefore become inaccurate for large distances *L*. For this reason, social insects also use guidance by local landmarks or so-called snapshots, which produce so-called local movement vectors [19]. Using this path-segmented navigation strategy reduces the error by global path integration. Previous studies on ants and bees have shown that the animal’s PI state can help in facilitating these view-based LVs [46, 53].

Inspired by these findings, we developed a reward-modulated learning rule for LVs following the same form as for GVs. Like in the previous case of global-vector guided navigation, the agent initially explores the environment randomly and acquires and expresses vector memories through experience. Insects have shown to be innately attracted to prominent landmarks or views of their surroundings [54, 55]. Thus here we implement such a fixed behavior in order to use visited landmarks as a signpost. The agent is attracted within a radius of 0.6 m to the landmark until it reaches as close as 0.05 m, where attraction is lost. At this distance to the landmark, we assume that the agent can detect a stored view represented by binary units

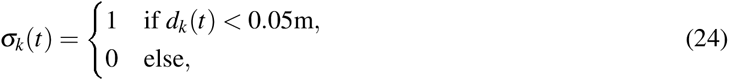

where *d*_*k*_(*t*) is the distance between the agent and the *k*th landmark. Active detection units elicit an eligibility trace *e*_*k*_, which is defined as

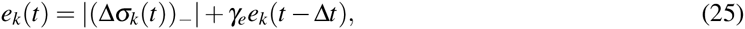

where *γ*_*e*_ = 0.995 is the discount factor of the decaying eligibility trace, and |(Δ*σ*_*k*_(*t*)) _–_| is the absolute negative change of *σ*_*k*_. Thus, eligibility traces are activated as the agent leaves the landmark detection area to avoid overlapping eligibility traces and rewards. Furthermore, we consider only the most recently active eligibility trace *ê*_k_ relevant for vector acquisition and expression. The eligibility traces serve as context-dependent units in the learning circuit (see Fig. 5). Similar to GVs, the context units activate LV representations in circular arrays. The activity of the circular arrays is defined as

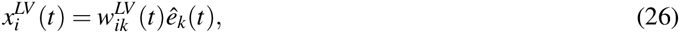

where 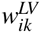 are the weights of the plastic synapses. These weights are modified using an update rule given by

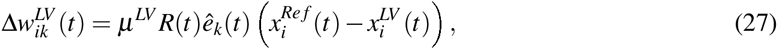

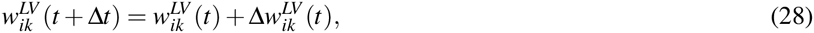

where *μ*^*LV*^ = 1 is the learning rate, and 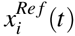 is a reference vector state in the direction 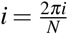, and *R*(*t*) is the total reward received. The reference vector state is simply a copied version of the PI mechanism, in which the memory layer is reset every time the agent detects a landmark. As such, the reference vector state represents a vector originating from the most recently visited landmark. In order to learn LVs from one landmark to another,the total reward *R*(*t*) does not only account for a food reward, but also for landmarks that lead to the feeder given by

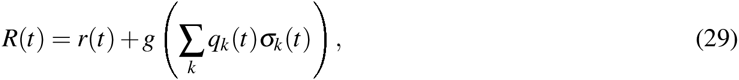

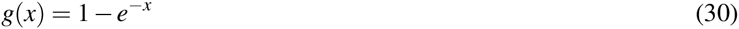

where *q*_*k*_ ≥ 0 are values assigned to the *k*th detected landmark, and *g* is a function that bounds landmark-evoked rewards to [0,1] for given non-negative arguments. Since the landmarks are initially unvalued (*q*_*k*_(0) = 0) to the agent, we apply a value system for learned LVs originating from respective landmarks, where each value is updated using

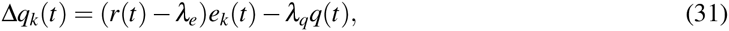

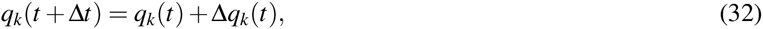

where *λ*_*e*_ = 10^‒5^ is an eligibility-based decay constant and *λ*_*q*_ = 10^‒6^ is a global decay constant. Thus, detected landmark values are discounted when the corresponding eligibility trace can not be associated with a delayed food reward *r*. Values also decay globally over time with constant *λ*_*q*_ to achieve competition between LVs. Note that, although we apply *K* landmark detection units with index *k* ϵ [0, *k* – 1], *k* does not have to correspond to the number of landmarks in the environment. Rather, LVs with a value below a certain threshold could be discarded for newly detected landmarks.

Finally, the output signal from the LV learning circuit is given by

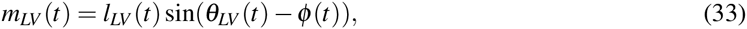

which follows a similar orientation error compensation as for the other vector representations. The angle *θ*_*LV*_ is computed from the position of the maximum activity in the circular array 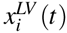 using the population vector average (see Eq. 10).

The final motor command ∑ for landmark-guided navigation (see Fig. 6) is given by the linear combination

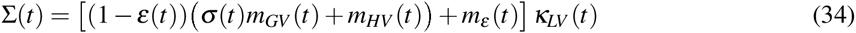

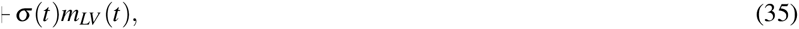

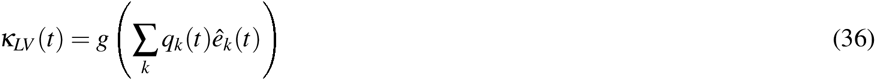

where random exploration *m*_*ε*_ and goal-directed navigation *m*_*GV*_ are inhibited by the time-varying signal *k*_*Lv*_ corresponding to the expression of rewarding LVs. Thus, an agent navigating through landmark-rich environments is biased towards the expression of LVs over GVs as observed in social insects [1, 15]. See S4 Text for a pseudocode description of our learning algorithm.

**Figure 6:**
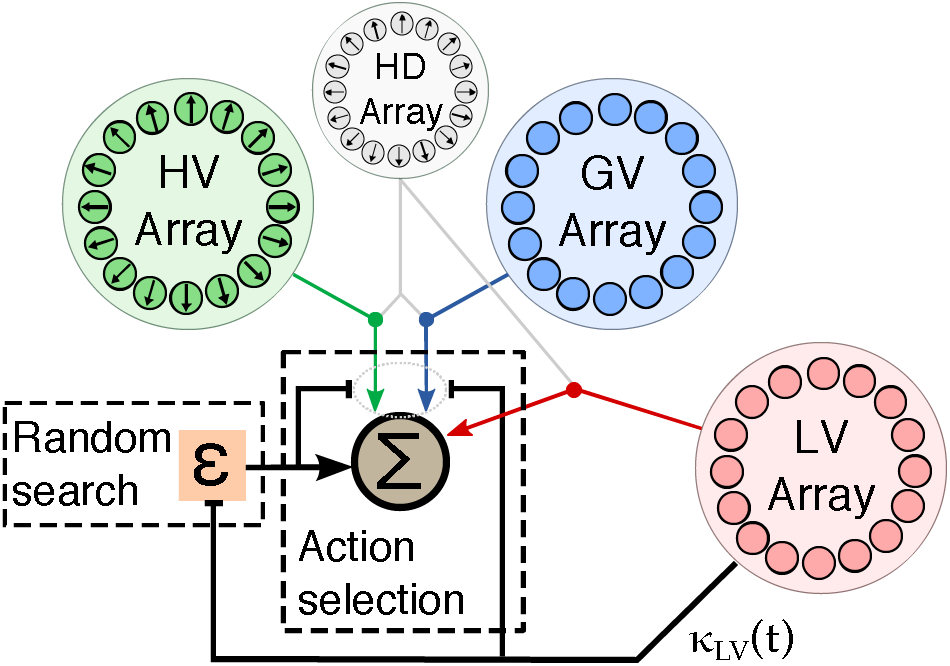
Combining global (GV) and local vector (LV) information for landmark-guided navigation. The expression of LVs surpresses random exploration and GV-guided navigation in the agent.

## Results

Using the proposed model embedded as a closed-loop control into a simulated embodied agent, we carried out several experiments to validate the performance and efficiency in navigating the agent through complex and noisy environments. We will further demonstrate that the generated behaviors not only resemble insect navigational strategies, but can actually predict observed behavioral parameters of social insects.

### Path integration (PI) in noisy environments

It has been shown, both theoretically and numerically, that PI is inherently prone to error accumulation [35, 56]. Studies have focused on analyzing resulting errors from using certain coordinate systems to perform PI [56–58]. Here we apply a system of geocentric static vectors (fixed preferred orientations) and analyze the effect of noise on the resulting error. How can noise be characterized in PI systems? Both artificial and biological systems operate under noisy conditions. Artificial systems, such as robots employ a multitude of sensors which provide noisy measurements, and generate motor outputs that are similarly noisy. Rounding errors in their control systems can be an additional source of noise. In animals, noise is mainly attributed to random influences on signal processing and transmission in the nervous system, including synaptic release and membrane conductance by ion channels and pumps (see [59] for review).

In order to validate the accuracy of the PI mechanism, we measure the positional errors of the estimated nest position with respect to the actual position over time. In the following experiments, we averaged positional errors over 1000 trials with trial duration *T* = 1000s (simulation time step Δ*t* = 0.1s). Fig. 7A shows the distribution of positional errors for three different sensory noise levels (1, 2, and 5 %). The distribution of errors follows a two-dimensional Gaussian distribution with mean 0.0 (nest) and width (*δ*_*r*_).

**Figure 7:**
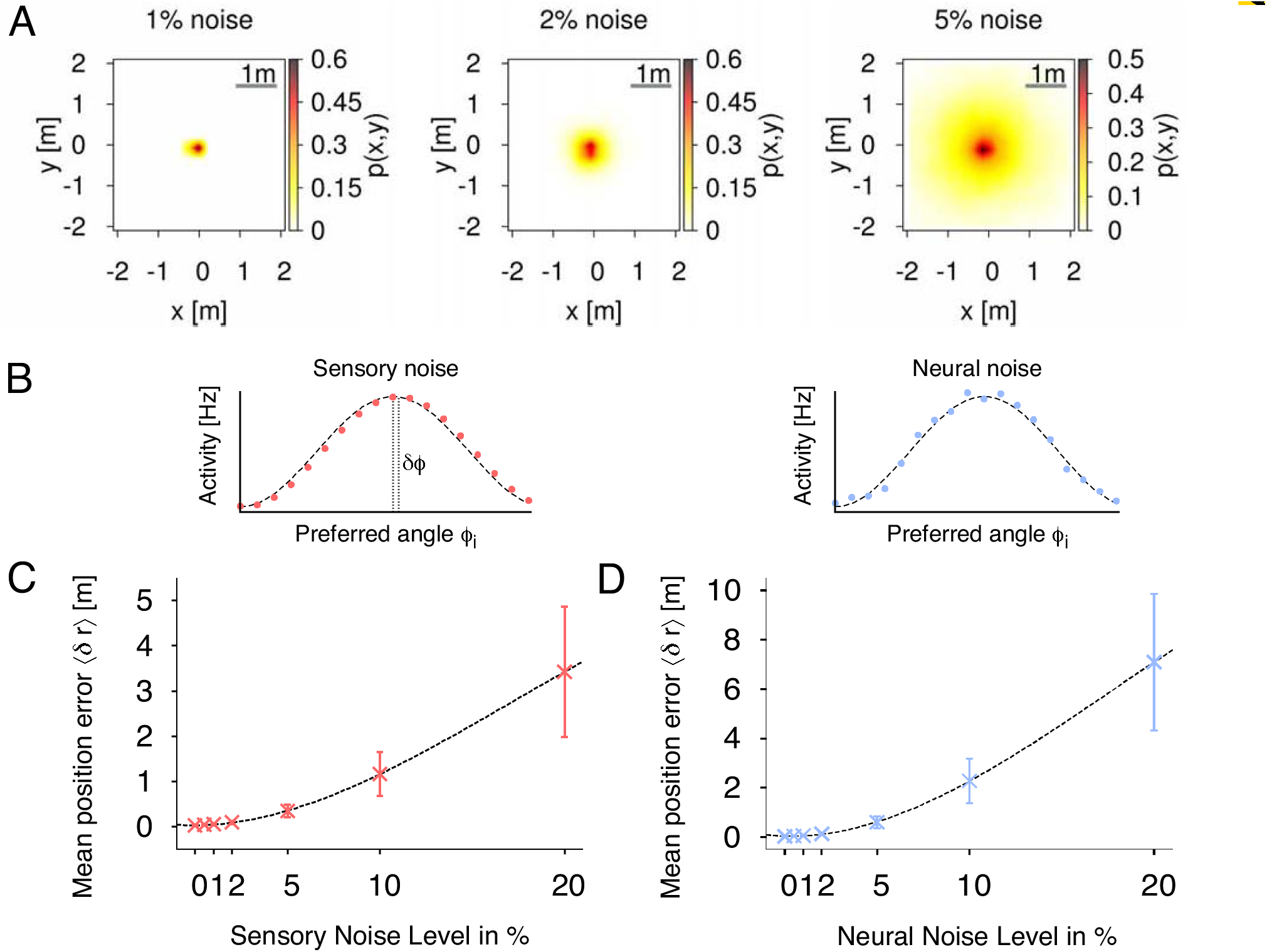
Path integration (PI) accuracy under the influence of external noise. A) We evaluate the accuracy of the proposed PI mechanism by using the mean positional error averaged over each time step during each trial. Distribution of positional errors for different sensory noise levels: 1%, 2%, and 5%. B) Examples of population-coded HD activities with correlated and uncorrelated noise. C) Mean position errors 〈*δ*_*r*_〉 (± S.D.) in PI with respect to fully correlated, sensory noise levels averaged over 1000 trials (fixed number of 18 neurons per layer). D) Mean position errors (*δ*_*r*_) (± S.D.) in PI with respect to uncorrelated, neural noise levels averaged over 1000 trials (fixed number of 18 neurons per layer).

In population coding, neural responses are characterized by correlated or uncorrelated noise ([60], see Fig. 7B for examples). In the uncorrelated case, fluctuations in one neuron are independent from fluctuations in the other neurons. Correlated noise is described by fluctuations which are similarly expressed across the population activity, and therefore leads to a shift of the observed peak activity. Here, we numerically analyze the effects of correlated and uncorrelated noise on the accuracy of the proposed PI mechanism. Correlated noise is here defined as a shift *δ*ϕ of the peak activity, i.e. fully correlated noise, such that the compass input to the PI mechanism is given by

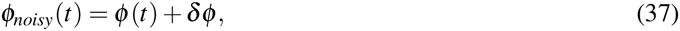

where *δϕ* is drawn from a Gaussian distribution *Ɲ*(0,2*πζ*_*sens*_) with sensory noise level *ζ*_*sens*_. Uncorrelated noise, also referred to as neural noise, is defined by adding fluctuations 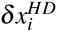 to the activities of the HD layer, which are drawn from a Gaussian distribution *Ɲ*(0, *ζ*_*neur*_) with neural noise level *ζ*_*neur*_.

Fig. 7C shows the effect of different degrees of sensory noise on the performance of PI for a fixed number of 18 neurons per layer averaged over 1000 trials. For noise levels up to 5% (equal to 18°), the observed mean position error increases only slowly and nonlinearly with values below 0.4 m demonstrating that our PI mechanism is robust for sensory noise up to these levels.

In Fig. 7D, we show mean position errors for different levels of uncorrelated noise. Similar to sensory noise, the errors first increase slowly and nonlinearly for noise up to 2%, while for noise larger than 5%, errors increase linearly. In comparison with sensory noise levels, uncorrelated noise leads to larger errors due to a more dispersed peak activity. However for noise levels up to 2%, mean position errors are well below 0.2 m indicating robustness of our PI mechanism with respect to uncorrelated noise.

In Fig. 8, we varied the number of neurons in the circular arrays of the PI mechanism for three different sensory noise level (0, 2, and 5 %). While the mean position error is significantly higher for 6 and 9 neurons, it achieves a minimal value for 18 neurons. For larger system sizes, the error only changes minimally. This is again mainly due to the coarse coding of heading directions. Interestingly, the ellipsoid body of the insect central complex contains neurons with 16-32 functional arborization columns (called wedges, see [61]). The numerical results here might point towards an explanation for this number, which efficiently minimizes the error.

**Figure 8:**
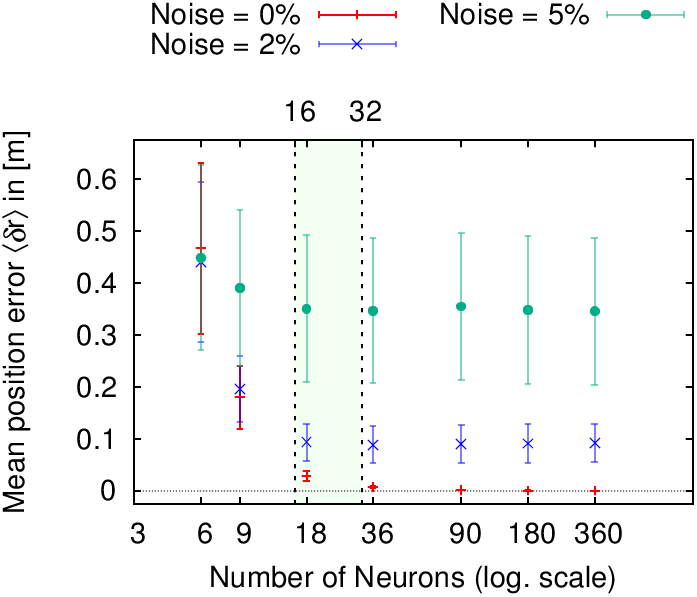
Mean positional errors 〈δ〉 (± S.D.) in path integration (PI) with respect to number of neurons per layer averaged over 1000 trials for three different sensory noise level (0, 2, and 5 %). In all three cases, the error reaches a minimum plateau between 16 and 32 neurons (colored area), which corresponds to the number of functional columns in the ellipsoid body of the insect central complex [61].

Besides errors resulting from random noise, there are also systematic errors observed in navigating animals. Both invertebrate and vertebrate species exhibit systematic errors in homing behavior after running an L-shaped outward journey (see [7] for review).[22] have examined such errors in desert ants by measuring the angular deviation with respect to the angle of the L-shaped course (see Fig. 9). In order to show that our mechanism is able to reproduce these errors, we fit our model against the desert ant data from Müller and Wehner [22] using the leaking rate *λ*(Eq. 7) of the PI memory layer as control variable. Using a leaking rate of *λ* ≈ 0.0075 resulted in angular errors most consistent with behavioral data. Leaky integration producing systematic errors is an idea that has been previously proposed [35, 62]. Thus, here our mechanism is not only performing accurately in the presence of random noise, but it also reproduces behavioral aspects observed in animals. In Table 1, we compare the accuracy and efficiency with other state-of-the-art PI models. Haferlach et al. [38] apply less neurons than our model, but we achieve a better performance in terms of positional accuracy with larger sensory noise. The model by Kim and Lee [39] applies 100 neurons leading to a fairly small positional error despite of 10% uncorrelated noise. Yet, both models do not account for random foraging as observed in insects. Desert ants were measured to freely forage average distances of 10-40 m depending on the species [63]. Indeed, if we reduce the foraging distance in our model, we achieve similarly small positional errors.

**Figure 9:**
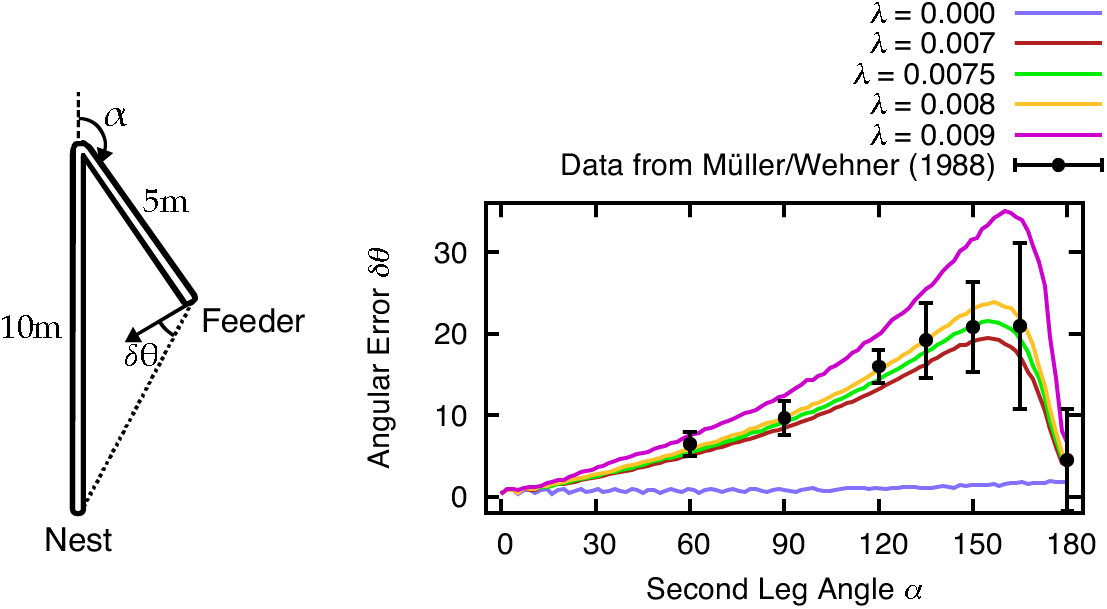
Systematic errors 56 of desert ant homing are reproduced by leaky integration of path segments. [22] tested the ants how accurate they return to the nest after following the two connected, straight channels with 10 m and 5 m length to the feeder (sketch modified from [22]). The second channel angle *α* was varied in 2.5° intervals for the simulation results. In our model, the leaking rate *λ* in the self-recurrent connections is used to fit the behavioral data [22]. We found that values *λ* ≈ 0.0075 accurately describe the observed systematic errors in desert ants.

**Table 1:** Comparison of existing path integration (PI) models in terms of accuracy and efficiency

| Model | Neurons | Noise level [%] | Positional error [m] | Foraging distance [m] |
| --- | --- | --- | --- | --- |
| Haferlach et al. [38] | 6 | 3 | $0.46 \pm 0.18$ | $\leq 5$ |
| Kim & Lee [39] | 100 | 10 | $0.020 \pm 0.002$ | $\leq 5$ |
| Our model | 18 | 5 | $0.351 \pm 0.140$ | $10 \pm 5$ |
| | 18 | 10 | $1.160 \pm 0.484$ | $10 \pm 5$ |
| | 18 | 5 | $0.070 \pm 0.037$ | $5 \pm 3$ |

### Global vector (GV) learning and goal-directed navigation

We introduced PI as a behavioral strategy to find the straight path back home. It also provides the animal with a spatial respresentation of locations, which can be used to form spatial memories [5]. Indeed, experiments have shown that desert ants are capable of forming such memories by using their path integrator [9, 64]. These memories are interpreted as so-called GVs, because the vector origin is fixed to the nest [19]. If the ant is forced to take a detour during a foraging trip, the deviation from the GV is compensated by comparing the GV with the current PI state [9].

In the previous section, we proposed a reward-modulated associative learning rule for GV learning. In order to test the performance of our insect-inspired model applying this learning rule, and to validate the use of learned vector representation in goal-directed navigation, we carried out several experiments under biologically realistic conditions. We apply the PI mechanism with *N* = 18 neurons per layer and a sensory noise level of 5%. In the first series of experiments, a single feeder is placed with a certain distance *L*_*feed*_ and angle *θ*_*feed*_ to the nest. The agent is initialized at the nest with a random orientation drawn from a uniform distribution on interval [0, 2*π*). In this naive condition, the agent starts to randomly search in the environment. If the agent is unsuccessful in locating the feeder after a fixed time *t*_*forage*_, it turns inward and performs homing behavior using only the PI mechanism. If the agent however finds the feeder, the current PI state is associated with the received reward, and stored in the weights to the GV array. The agent returns back home after the accumulated reward surpasses a fixed threshold. Each trial lasts a fixed maximum time of 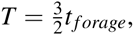, before the agent is reset to the nest position. On subsequent foraging trips, the agent applies the learned vector representation and navigates along the GV, because the exploration rate is decreased due to the previous reward. If the agent finds the feeder repeatedly, the learned GV stabilizes and the exploration rate decreases further.

Fig. 10 demonstrates two of such experiments for a feeder with a distance of *L*_*feed*_ = 2 m and angle *θ*_*feed*_ = 180°, and with a distance of *L*_*feed*_ = 10 m and angle *θ*_*feed*_ = 90°, respectively, from the nest. In Fig. 10A, we show the trajectories of the agent during five trials for the feeder placed 2 m away from the nest. The trial numbers are color-coded (see colorbox). During the first trial, the agent has not visited the feeder yet and returns home after *t*_*forage*_ = 400 s of random search. During the second trial (see yellow-colored trajectory), the agent finds the feeder and learns the GV representation from the PI state (see 10C). Here the red dotted line indicates the correct angle *θ*_*feed*_ = 180° to the feeder, while the cyan-colored line is the average angle estimated from the synaptic strengths. After the agent reaches the feeder the first time, the learned vector is visibly inaccurate with respect to the correct feeder angle. Furthermore, as the exploration rate has only decreased by a small amount (Fig. 10E), the subsequent outward trip of the third trial (green-colored trajectory) is still significantly controlled by the random term. This allows for compensating initial inaccuracies in learning due to PI errors. Because the agent reaches the feeder a second time, the average angle is closer to the feeder angle. The repeated visits to the feeder decrease the exploration rate due to the received reward (red line). In the final two trials, the agent navigates to the feeder on a stable trajectory (i.e., low exploration rate) demonstrating that the learning rule is robust for goal-directed navigation in noisy environments.

**Figure 10:**
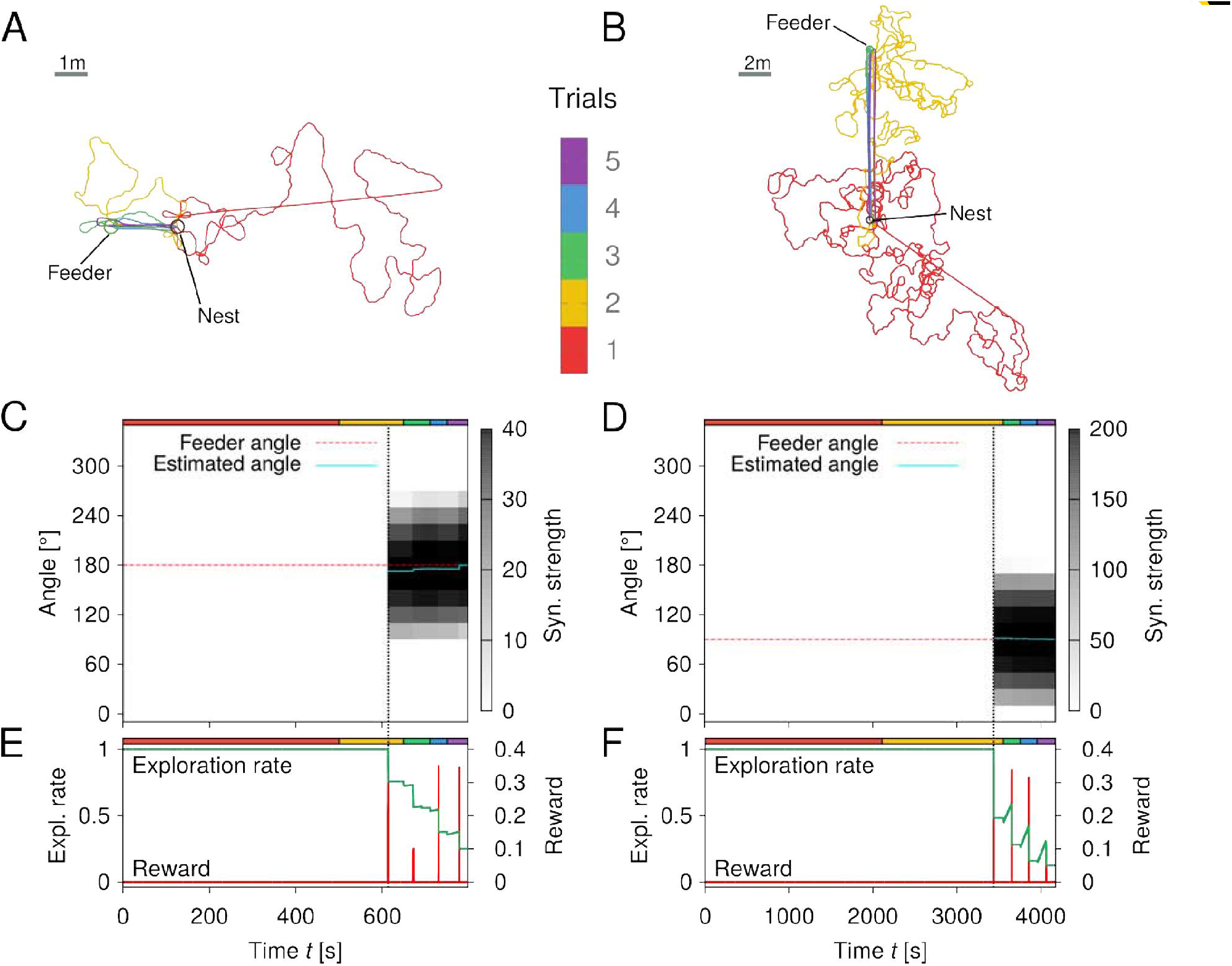
Learning walks of the simulated agent for a feeder placed *L*_*feed*_ = 2 m and *L*_*feed*_ = 10 m away from the nest, respectively. Trajectories of the agent for five trials with a feeder in A) 2 m distance and 180° angle to the nest, and B) 10 m distance and 90° angle to the nest. Each trial number is color coded (see colorbar). Inward runs are characterized by straight paths controlled only by PI. See text for details. Synaptic strengths of the GV array changes due to learning over time (of the five trials) for the feeder of C) 2 m distance and D) 10 m distance. The estimated angle *θ*_GV_ (cyan-colored solid line) to the feeder is given by the position of the maximum synaptic strength. Exploration rates and food reward signals with respect to time for the feeder of E) 2 m distance and F) 10 m distance.

This robustness and stability of the learning rule holds also for larger scales. We increased the distance of the feeder to the nest to *L*_*feed*_ = 10 m and rotated the angle to *θ*_*feed*_ = 90°. Fig. 10B demonstrates the learning walks of the agent. Here we increased the foraging time of the agent to *t*_*forage*_ = 2000 s. In this experiment, the agent finds the feeder during the second trial. In doing so, the agent is able to acquire an accurate vector representation (Fig. 10D) resulting in stable trajectories towards the goal for the final three trials, which is again due to a low exploration rate (Fig. 10F). Note, that the represented GV lengths are characterized by a linear relationship, as the maximum synaptic strength for the 10 m feeder is five times larger than the one for 2 m.

A single-feeder environment, however, does not represent realistic environments encountered by social insects. Therefore, the next experiment contains an environment with randomly placed feeders. 50 feeders are placed uniformly in a circular radius of 40 m with mininum distance of 3 m between them. Like in the first experiment, we apply *N* = 18 neurons in the circular arrays and a sensory noise level of 5%. After each of 1000 trials with foraging time *t*_*forage*_ = 1000 s, we measured the exploration rate, homing success, and goal success. The homing success is determined by whether the agent reaches the nest during the given total time *T*. Goal success is defined as, whether or not the agent has received any reward during a trial. We processed homing and goal success as a running average with respect to trials. Fig. 11 shows the results of the experiment in a randomly generated environment. In the top row, we show two-dimensional density plots of the agent’s trajectories before and after learning. The color-coded probabilities *p*(*x*,*y*) were measured by counting each trajectory point within squared boxes with a spatial resolution of 0.2 m and normalizing by the number of points in each box, respectively. The agent initially performs random search in the environment. When the agent encounters a feeder (see green-colored circle), the exploration rate decreases. After learning, i.e., for low exploration rates *ε*(*t*), the agent performs goal-directed navigation to the learned feeder. This is further indicated by the bottom graph, which shows exploration rates after each trial, as well as running averages of homing and goal success, respectively. When the GV is learned and the exploration rate is sufficiently low, the goal success rate approaches the value one for large number of trials. Thus, GV learning and goal-directed navigation is shown to be stable and robust in a realistic, randomly generated environment.

**Figure 11:**
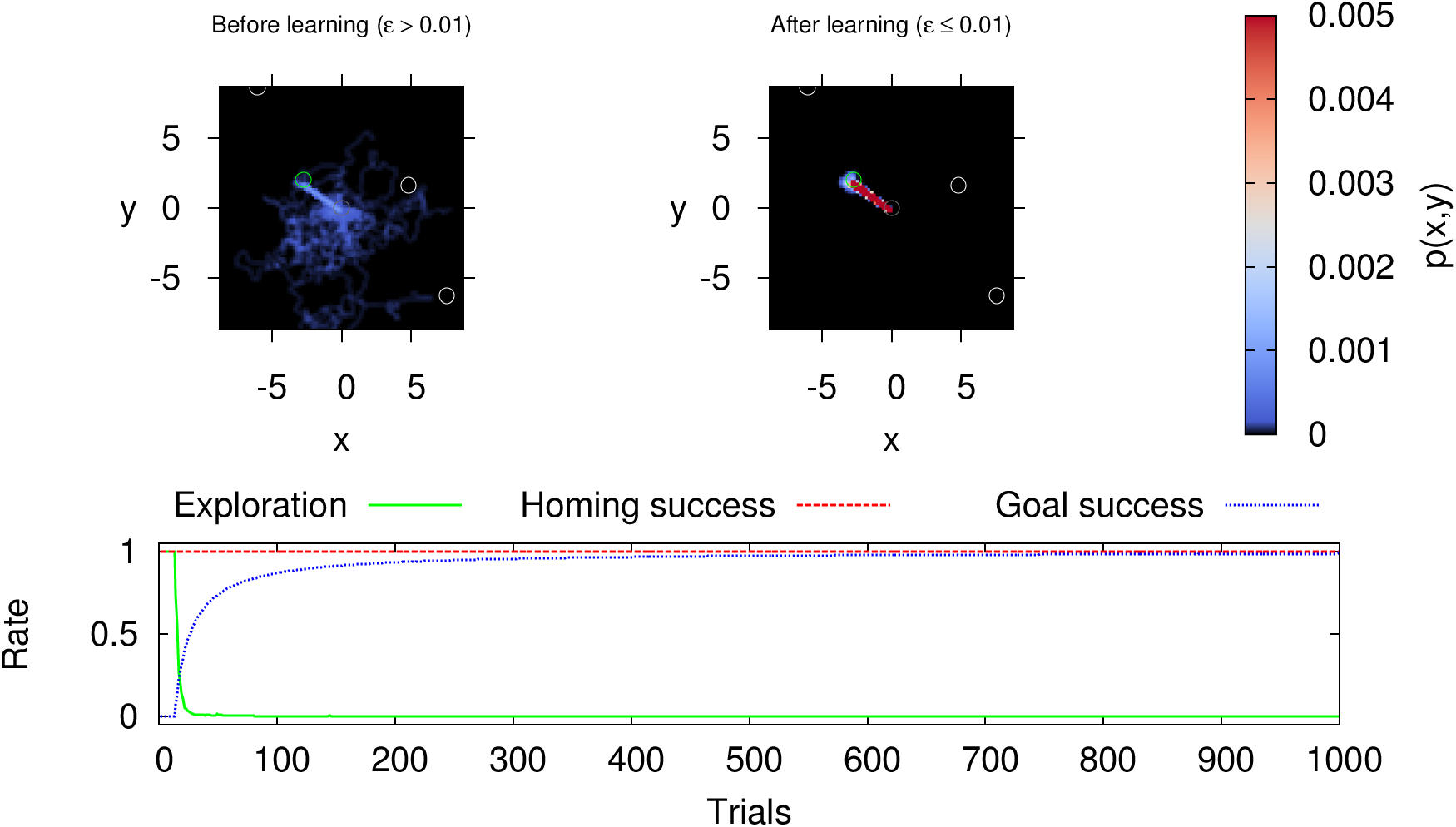
Global vector (GV) learning and goal-directed navigation in an environment with randomly placed feeders. Top row: Two-dimensional density plots were measured by counting trajectory points within squared boxes of spatial resolution of 0.2 m, before being normalized by the number of points within the box. The resulting probabilities *p*(*x*,*y*) are color-coded, where black corresponds to zero probability. Bottom row: Exploration rate after each trial, and running averages of homing and goal success with respect to trials. See text for details.

In Fig. 12, we simulated 100 learning cycles with different randomly generated environments, each consisting of 100 consecutive trials. In Fig. 12A, we show the mean exploration rate, and the running averages of mean homing and goal success rates with respect to trials (foraging time *t*_*forage*_ = 1000 s, averaged over 100 cycles). During the 100 trials, learning converges on average within the first 20 trials given by a low mean exploration rate. Like in the previous experiment, the agent reaches the feeder in every trial after convergence is achieved. This is indicated by the goal success approaching one. Average homing success is one for every trial,which results from sufficient searching behavior and the given total time *T*. The convergence of the learning process is dependent on the foraging time, because longer time allow for longer foraging distances, and thus larger search areas. Therefore, we varied the foraging time *t*_*forage*_ = 200,400,600,800, and 1000 s and measure the mean goal success rate after 100 trials averaged over 100 cycles (Fig. 12B). The results indicate that for longer foraging times, the mean goal success rate approaches one and its variance decreases. However, by measuring the averaged ratio of learned vector and nearest feeder distance, we show that this ratio decreases for larger foraging times (Fig. 12C). Thus, there is a trade-off with respect to convergence and reward maximization, leading to an optimal foraging time. Desert ants have been shown to increase their foraging times up to a certain value, after which it saturates [13]. This adaptation of foraging time might be indicated by the trade-off resulting from our model.

**Figure 12:**
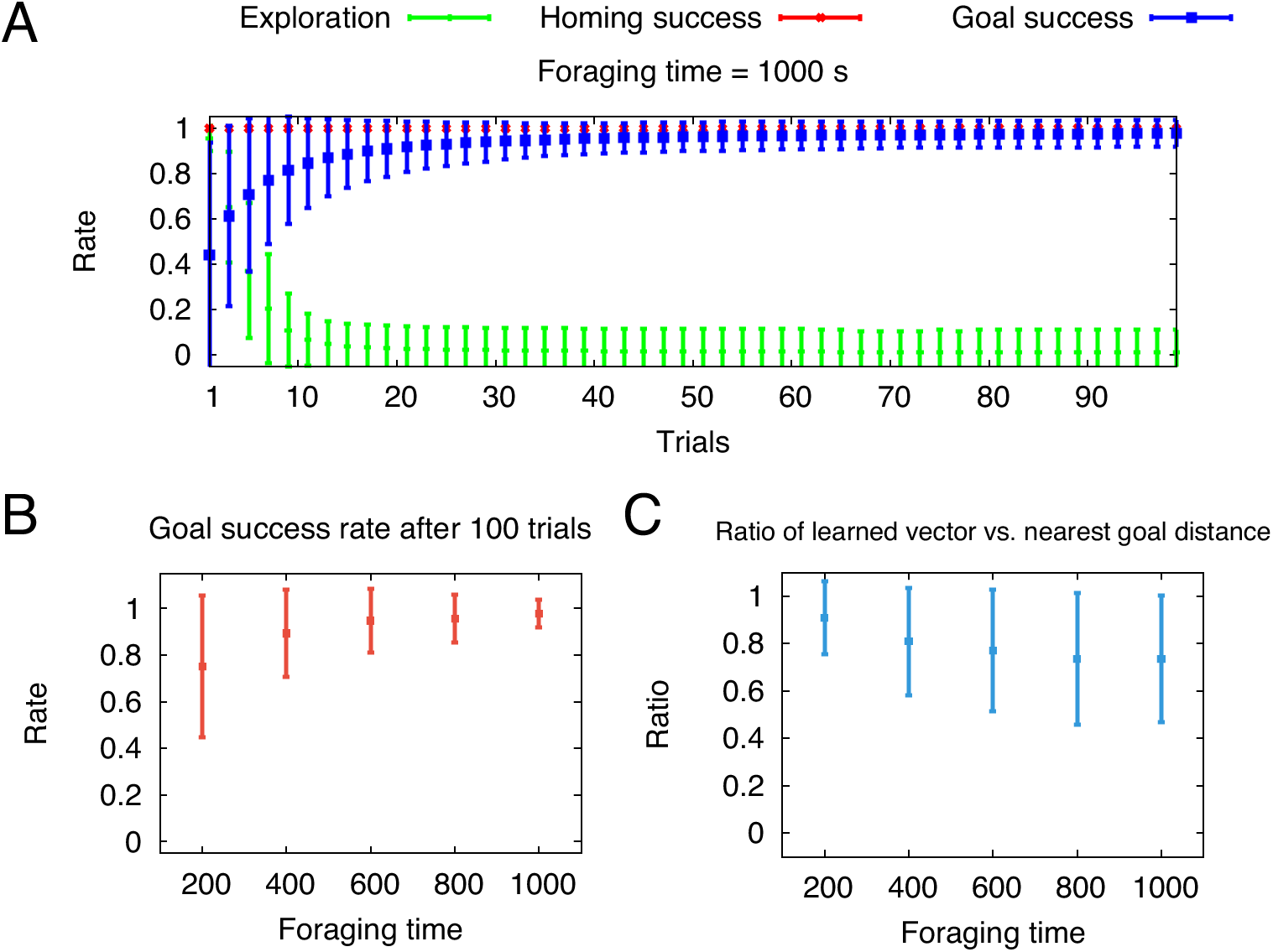
Longer foraging durations during global vector (GV) learning increase the average goal success rate, but decrease the ratio of learned global vector and nearest feeder distance. A) Mean exploration rate and running mean goal success and homing rate (± S.D.) with respect to trials averaged over 100 cycles of randomly generated environments (foraging time *t*_*forage*_ = 1000 s). Goal success is defined by whether a feeder was visited per trial. The homing rate is determined by the agent’s return to the nest within the given total trial duration *T*. *B*) Mean goal success rate after 100 trials with respect to foraging time *t*_*forage*_ averaged over 100 cycles. C) Mean ratio of learned GV distance and nearest feeder distance with respect to foraging time *t*_*forage*_ averaged over 100 cycles.

### Local vector (LV) learning and route formation

Social insects have been shown to be guided by so-called LV memories which encode sufficient information about both the direction and distance of a route segment [18]. Its acquisition and expression is supported by PI, as well as local and panoramic views [15]. Here we present a neural mechanism to acquire a population-coded representation of LV memories stored in the synaptic weights connecting to a circular array of neurons. LVs are expressed when the agent reaches a prominent landmark.

In order to demonstrate the learning dynamics of the LV learning, we generated an environment with two additional landmarks at (*x*_*LM*1_ = 1 m,*y*_*LM*1_ = 2 m) and (*xLm*_2_ = 1 m,*y*_*LM*2_ = 4 m), respectively, and a feeder at (*x*_*feed*_ = 0 m,*y*_*feed*_ = 5 m). Note, that the nest also serves as a landmark. The agent now forages for 1000 trials with foraging time *t*_*forage*_ = 150 s. Fig. 13 shows the trajectories of the agent during different trial intervals. During the first 120 trials, the agent randomly searches in the environment. When the agent reaches the feeder, LVs are acquired and expressed in subsequent trials (see trials 121-150). For the trials 151-180, random exploration further decreases as the LV from the nest to the first landmark is learned. Finally, every route segment is robustly expressed by LVs and a stable route is thus formed for the trials 181-1000. We emphasize that the order of landmarks that the agent walks by is not predefined, but rather learned by association of the landmark eligibility traces with a reward signal. Since initially there is only reward given at the feeder, LV memories are acquired recursively from the feeder towards the nest.

**Figure 13:**
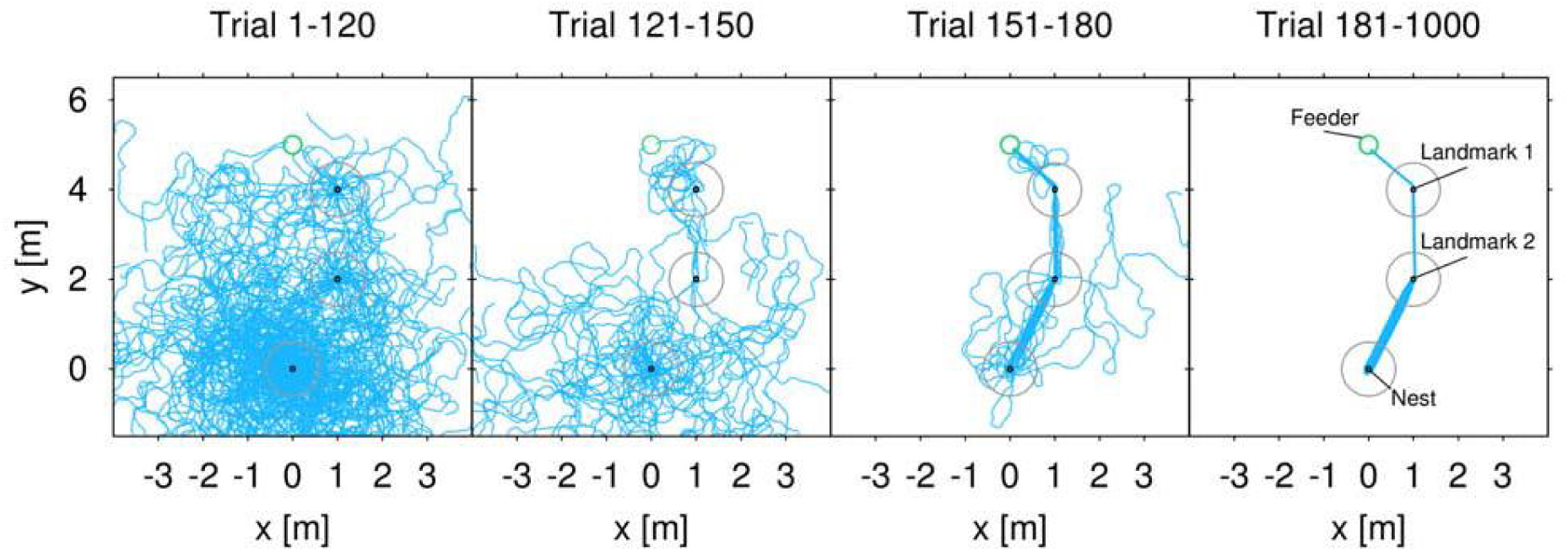
Route formation through navigating with local vectors (LVs). The environment is composed of two landmarks (black circles) at (*x*_*LM*1_ = 1 m, *y*_*LM*1_ = 2 m) and (*x*_*LM* 2_ = 1 m, *y*_*LM*2_ = 4 m), and a feeder (green circle) at (*x*_*feed*_ = 0 m, *y*_*feed*_ = 5 m). The nest (black circle at origin) serves as an additional landmark. Landmarks are attractive to the agent when approaching the landmark within a radius of 0.6 m (see gray circle). See labels in the outer right panel. Trials 1-120: The agent randomly explores the environment, 121-150: First transition as the second LV (landmark 1 to landmark 2) is robustly expressed, 151-180: Second transition as first and final LV is robust expressed, 181-1000: Random exploration approaches zero, thus LV learning converges and the agent forms a stable route.

## Discussion

The central question of this paper is how vector memories as observed in behavioral studies on bees and ants [9, 11, 18, 19] are encoded in insect neural systems, and how such representations are stored and retrieved in a biologically plausible way. Here, we have shown that a computational model based on population-coded vector representations can generate efficient and insect-like navigational behaviors in embodied agents. These representations are computed and stored using a simple neural network model combined with reward-modulated associative learning rules. Thus, the proposed model is not only accounting for behavioral aspects of insect navigation, but it further provides insights in possible neural mechanisms in relevant insect brain areas, such as the central complex. In the following, we will discuss certain aspects of our model with respect to neurobiological findings in insects. Furthermore, we provide comparisons to other state-of-the-art models of vector-guided navigation [33, 41].

### Head-direction (HD) cells & path integration (PI)

A main property of the PI mechanism of our model is that it receives input from a population of neurons, which encode for allothetic compass cues. Here, we apply a cosine response curve for coarse encoding of orientations. Such a mechanism was previously applied by other models [38, 39]. Neurons in the central complex of locusts contain a population-coded representation of allothetic compass cues based on the skylight polariza-tion pattern [30]. Similarly, central complex neurons in the Drosophila brain encode for heading orientations based on idiothetic self-motion and visual landmarks. Seelig and Jayaraman [23] measured the fluorescent activity of genetically expressed calcium sensors indicating action potentials, while the fly was tethered on an air-suspended track ball system connected to a panoramic LED display. Any rotation of the fly on the ball is detected and fed back by corresponding motions of the visual scene on the display. The activity of 16 columnar neurons, which display the full circular range, generates a single maximum, which moves according to the turns of the fly on the ball. Interestingly, even though the representation is generated by visual stimuli, it can be accurately maintain solely by self-motion cues over the course of several seconds in the dark. A recent study on dung beetles [65], which navigate completely unaffected by landmarks, has shown that celestial compass cues are encoded in the central complex revealed by electrophysiological recordings. Taken together, it is likely that the central complex of social insects contains a similar neural coding of polarization‐ and landmark-based compass cues. Not only is the central complex function and anatomy highly conserved across insect species, but behavioral experiments on ants and bees also suggest the central role of using polarization and landmark cues for navigation. Our model further predicts allothetic and landmark-based compass cues to be involved in global and local PI mechanisms. Such neural computations have yet to be observed in experiments, ideally by applying the tethered track ball setup described in [23]. In our model, we assume that the agent’s walking speed is neurally encoded as a linear signal that modulates the amplitude of HD activities by an additive gain. A similar, so-called gater mechanism has been applied in a model by Bernardet et al. [66]. Such linear speed signals have recently been found to be encoded by neurons in the rat’s medial entorhinal cortex [67] as well as the cockroach central complex [68]. This shared encoding mechanism indicates the necessity of linear velocity components for accurate PI [69]. The temporal accumulation of speed-modulated HD signals in our model is achieved by a self-recurrent connection. Biologically, these recurrent connections can be interpreted as positive feedback within a group of neurons with the same preferred direction. Since our model applies PI as a scaffold for spatial learning, we apply this simplified accumulation mechanism to avoid random drifts observed in more complex attractor networks [70], which were applied in previous PI models [36, 71]. We were also able to test the leaky-integrator hypothesis [62] by fitting a single leakage parameter to observed behavioral data from desert ants [22]. The leakage parameter decreases the self-recurrent connection weight for leaky integration. A HV representation is computed by using a cosine weight kernel, which was also used in [66]. Such a connectivity acts on each represented direction by adding the projections from other directions, respectively. This leads to the formation of an activity pattern with a single maximum across the population. The angle of the represented vector is readout by averaging the population vectors, while the distance is encoded by the amplitude of the population activity. We show that such a readout of a population-coded vector is sufficient to generate robust homing behavior in an embodied agent. Furthermore, it allows for accurate localization required for spatial learning of locations.

Our extensive numerical analysis of noise affecting PI accuracy has shown two predictions. First, PI accuracy seems to follow a similar function with respect to the noise levels for both the fully correlated and uncorrelated random fluctuations. While uncorrelated noise could be further filtered depending on the system size *N*, decorrelation of sensory input noise could be achieved by adding inhibitory feedback as shown in a model by [72]. Second, we varied the number of neurons *N* per layer for different levels of fully correlated noise, which predicts an accuracy plateau between 16 and 32 neurons where the accuracy will not increase for larger systems. This indicates that such a number of partitions for representing orientation variables is efficient and accurate enough. Interestingly, most prominent neuropils of the central complex exhibit a similar number of functional columns [61]. The central complex has been shown to be involved in sky compass processing [30], spatial orientation [23], and spatio-visual memory [73, 74]. Its columnar and reverberating connectivity further supports the functional role of integrating orientation stimuli. These evidences suggest that the proposed circular arrays representing navigation vectors might be encoded in the central complex. We conclude that further experiments are needed to unravel how PI is exactly performed in the insect brain by closely linking neural activity and circuitry to behavioral function.

### Reward-modulated vector acquisition and the role of motivational context

PI provides a possible mechanism for self-localization. As such, it has been shown experimentally that social insects apply this mechanism as a scaffold for spatial learning and memory [5]. Here we propose a reward-modulated associative learning rule [43–45] for acquiring and storing vector representations. The acquisition and expression of such vector memories depend on the context during navigation. For GVs, the context is determined by the foraging state, which we model as a binary unit. Indeed, behavioral studies on desert [75] and wood ants [76] have shown that expression of spatial memories is controlled by an internal state in a binary fashion. The association of the context with a reward signal, received at the feeder, drives synaptic weight changes corresponding to the difference between the current PI state and the respective weight. As this difference is minimized, the weights converge towards values representing the PI state when the reward was received at the feeder. Thus, like the HV, GVs are population-encoded with the angle determined by the position of the maximum activity and the length determined by the amplitude of the activity. To our knowledge, this is the first model that applies such a neural representation to perform vector-guided navigation. Previous models, such as [33, 41], do not provide possible underlying neural implementations of the PI-based stored information used for navigation. The HD accumulator model [33] argued that vector information is stored in so-called shortcut matrices, which are subsequently used for navigating towards goals. Similarly, the Cruse and Wehner model [41] stored HVs as geocentric coordinates in an address-based memory. While both models offer sufficient mechanisms in order to generate vector-guided navigation, they neither seem biologically plausible nor provide any explanations how such information is dynamically learned during navigation.

Our proposed encoding of GVs is validated by recent findings from a behavioral study on wood ants [76]. The authors carried out a series of novel experimental paradigms involving training and testing channels. In the training channel, ants were trained to walk from their nest to a feeder at a certain distance, before they were transferred to the testing channel. There, they measured the expression of vector memories by observing the behavior. The authors showed that vector memories are expressed by successful association of direction and distance, therefore such memories might be encoded in a common neural population of the insect brain. The acquisition of vectors were rapid after 4-5 training trials, which corresponds to the rapid vector learning shown by our model during learning walks (Fig. 10). However, the authors mainly examined the expression of homeward vector memories, which have not been shown in our model. Instead, we argue that PI alone can guide the agent back to the nest.

Two major higher brain areas in social insects exhibit experience-dependent plasticity due to foraging activity: the mushroom bodies [77] and the central complex [78]. The mushroom bodies are paired neuropils known to be involved in olfactory learning and memory [79], as well as visual learning in discrimination tasks [80]. Studies on the central complex across various insect species have revealed its role in visual object localization [81] and visual learning [82], motor adaptation [83], spatio-visual memory [23, 73, 74], as well as polarization-based compass [30]. A common coding principle in the central complex appears to be the topological mapping of stimuli within the full azimuthal circle [25]. Both higher brain neuropils involve the functional diversity of multiple neuropeptides and neurotransmitters [84]. The short neuropeptide F is a likely candidate influencing the foraging state, as it has been shown to regulate feeding behavior and foraging activity after starvation [85]. Based on this evidence, we conclude that the population-coded vector memories described by our model are likely to be found in the central complex. Nonetheless, we do not exclude the possibility of possible interactions between the central complex and the mushroom bodies involved in spatial learning and navigation, which is supported by recent findings on novelty choice behavior in Drosophila [86].

### Local vector (LV) learning and route-guided navigation

Our model suggests a critical role of PI for route-guided navigation in helping the acquisition of so-called LVs. Segmenting a GV into a route of shorter LVs is useful, because it reduces distance-based, accumulated PI errors. Evidence for our hypothesis is based on several behavioral studies on ants [15, 18, 19, 53] as well as bees [87, 88]. LV memories are not just fixed sensori-motor mappings, but rather learned allothetic directions[19]. During navigation in cluttered environments, insects are attracted towards prominent landmarks in the near surroundings [54, 55]. The expression of LVs is exclusively associated with such local views [15]. The encoding of LV directions are primarily based on the skyline panoramic view [16] as well as the polarization compass [89]. Recent experiments on desert ants suggest that odometry is applied to gauge and learn distances of LVs [18]. These behavioral findings are all in line with our model for LV learning. We further hypothesize that, like for GV memories [76], LVs encode both direction and distance in a common neural population.

In our model, the acquisition and expression of LV memories depend on detection events of local landmarks. Such detection events elicit an eligibility trace, which is associated with the reward received at the subsequent landmark or feeder. Experiments have shown that desert ants exhibit short-term memories of visual experiences that persist over a long period of about 30-100 s [90]. Our model performs local PI based on the most recently visited landmark to learn LVs. The compass cues used as input to local PI could also easily be inferred by a purely visual compass based on the surrounding panorama [91]. Using this model for LV learning, we have shown that the agent is able to learn to navigate on a stable route in the presence of external sensory noise. The Cruse and Wehner model [41] also presents such a landmark-guided navigation of foodward and homeward routes, but their model had LVs preset in static memory banks. Thus, their model does not explain how such vector memories can be acquired during foraging. Furthermore, their model assumes a fixed-length LV, which has shown to be unlikely in experiments with desert ants [18]. Although our model does not present route navigation towards the nest, it can easily be extended to learn both based on the foraging state. Finally, we see that our model has the capacity to work in parallel with view-based navigation models [92, 93]. A recent model [93] applied a spiking neural network based on mushroom bodies circuitry to achieve visually based route navigation. Their model suggests that associative learning and sparse coding in the mushroom bodies allows for extracting information from up to hundreds of visual scenes. We conclude that the interactions of mushroom bodies and central complex orchestrate the complex behavior observed in navigating social insects. A hierarchical structure of learning mechanisms acting on different timescales leads to robust navigational strategies [1]. GV learning is a strategy which is flexible and rapid enough to allow revisiting goals after a few successful trials. This flexibility allows to refine the subsequent travels by route segments first as PI-based LVs, later as purely view-based image matching leading to stereotyped and efficient routes.

## Conclusion

We proposed a novel computational model for PI and the acquisition and expression of vector memories in simulated, embodied agents. Other than previous models [33, 41], we provide plausible neural implementations of the underlying control and learning mechanisms. Our model has shown to be sufficient in generating robust and efficient behaviors, which have been observed in social insect navigation. Besides behavioral observations, our model also provides predictions about underlying neural implentations regarding structure and plasticity of related neural circuits in the insect brain [94]. We discussed our findings in the context of neurobiological evidences related to two higher brain areas of insects, the central complex and the mushroom bodies. We therefore conclude that our model offers a novel computational approach for studying vector-guided navigation in social insects, which combines neural mechanisms with their generated behaviors. This can guide future behavioral and neurobiological experiments needed to evaluate our findings.

## Acknowledgments

We thank Florentin Wörgötter at the Department of Computational Neuroscience in Göttingen, where most of this work was conducted. DG & SD thank Taro Toyoizumi and his lab members at RIKEN BSI for fruitful discussions. We thank James Humble for comments on the manuscript.

## Contributions

Conceived and designed the experiments: DG SD PM. Performed the experiments: DG. Analyzed the data:DG SD PM. Contributed reagents/materials/analysis tools: DG SD. Wrote the paper: DG SD PM.

## Supporting Information

**S1 Table**

**Model variables & parameters.** Further information on variables and parameters used throughout this work.

**S1 Text**

**Experimental platforms & foraging statistics.** Further information on experimental platforms used throughout this work.

**S2 Text**

**Directed walk using sinusoidal phasors.** Here we derive how the agent performs directed walk by using sinusoidal phasors. We show that such a description leads to stable dynamics, which drives the actual heading orientation towards the desired orientation. It further allows for vector substraction as presented in our proof.

**S3 Text**

**Deriving a gradient-based learning rule for *β*.** Here we derive a gradient-based learning rule for the parameter *β*, which determines how quickly the exploration rate of the agent converges to zero based on experienced reward.

**S4 Text**

**Pseudocode.** Pseudocode description of our learning algorithm for vector-guided navigation.

